# Lymph-Circulating Tumor Cells show distinct properties to Blood-Circulating Tumor Cells and constitute extraordinary efficient metastatic precursors

**DOI:** 10.1101/444091

**Authors:** Sulma I Mohammed, Odalys Torres-Luquis, Elwood Walls, Frank Lloyd

## Abstract

The molecular properties of tumor cells as they exit the primary tumor into the afferent lymphatics en route to the sentinel lymph nodes (SLNs) are not yet known. We developed an innovative technique that enables the collection of lymph and lymph-circulating tumor cells (LCTCs) en route to the SLN in immunocompetent animal model of breast cancer metastasis. We found that LCTCs and blood circulating tumor cells (BCTCs) as exited the primary tumor shared similar gene and protein expression profiles that were distinct from those of primary tumors and lymph node metastases (LNMs) despite their common parental cell origin. LCTCs but not BCTC exist in clusters, display a hybrid epithelial/mesenchymal phenotype and cancer stem cell-like properties and constitute extraordinarily efficient metastatic precursors. These results demonstrate that tumor cell metastasizing through the lymphatic are different from those spread by the blood circulation. The contribution of these cells to overall peripheral blood CTC is important in cancer therapy. Whether these two types of cells occur in cancer patients remain to be determined.

**Statement of significance:** The presence of tumor cells in the SLN denotes poor prognosis and worse patient outcomes. We have developed a capability to routinely collect these tumor cells before they reach their first stop, SLN, in their metastatic journey to distant sites. Examination of these cells and their lymph microenvironment revealed that they are molecularly different from their tumor of origin, their LNMs and their counterpart cells in the blood. This is the first time these cells are captured and studied. The approach will provide a new level of information that is highly relevant to our understanding of metastasis.

## INTRODUCTION

Breast cancer is the most common malignancy in women. The leading cause of breast cancer-associated death is metastasis (1). Although advances in early diagnosis and systemic adjuvant therapy targeting primary tumors have significantly improved survival in women with breast cancer, treatments for metastatic disease remain less effective. The problem in identifying therapies targeting metastatic disease is our incomplete understanding of tumor biology during the metastatic process. During metastasis, tumor cells detach from the primary tumor and may intravasate into and disseminate through the blood circulation or lymphatic system; either route of dissemination can lead to the venous circulation, as the lymphatics drain into the blood (2). In 80% of solid tumors, metastasis via the lymphatic system precedes metastasis via the vascular system. In many of these tumors, the lymph nodes are the first organ to develop metastasis. As a result, the tumor-draining lymph node, which is the sentinel lymph node (SLN), is accepted universally as the most powerful prognostic tool available for early-stage breast cancer and is often used in disease management (3)

Despite the clinical implications of tumor cell lymphatic spread and lymph node metastasis in breast cancer patient care and management, little is known about the cellular and molecular communication that takes place between the primary tumor and the sentinel node. In addition, lymphatically disseminated tumor cells in transit from the primary tumor to the local lymph node have never been characterized and compared to blood-borne tumor cells in the same host. Several studies have examined tumor cells discharged into the tumor venous drainage (4), but to our knowledge, there have been no experimental studies of LCTCs in transit from the primary tumor to the local draining SLN. The major reasons for this lack of knowledge have been the microscopic size of the afferent lymphatic vessels, the fragile nature of these vessels, the loss of pressure that occurs as soon as the vessels are punctured, and the difficulty in identifying and cannulating the lymphatic vessels en route to the SLN (5). The characterization of LCTCs and BCTCs may provide important information about the cascade of metastatic events.

Recently, accumulating evidence suggested that the microenvironment of the SLN is greatly influenced at a distance by the primary tumor, which secretes factors such as cytokines, exosomes, or enzymes that precondition the lymph node microenvironment, making the lymph nodes supportive metastatic niches for disseminating tumor cells (soil and seed hypothesis) (6,7). According to this understanding, the lymphatic fluid draining a primary tumor is expected to be rich in these premetastatic conditioning materials, which can serve as discriminating indicators of the tumor metastatic potential. The identification and monitoring of these premetastatic niche-inducing materials in situ in lymph draining a primary tumor can provide insights about immune recognition or immune priming in the SLN that are highly relevant to tumor treatment.

Here, we developed a unique microsurgical technique to collect lymph draining from a primary tumor. We have used an approach that is routinely practiced for the identification and mapping of the draining lymph nodes during the SLN dissection procedure in women diagnosed with breast cancer. The SLN concept implies that the tumor cells migrating from a primary tumor metastasize to a single lead draining node in the relevant lymph node basin (8). The injection of lymphazurin in the breast tissue around the area of the tumor permits the identification of one or more SLNs in the majority of patients. Taking advantage of this concept, we developed a technique to intercept the migration of tumor cells from the primary tumor to the SLN and collect both the lymph and the tumor cells therein. We collected a large enough volume of afferent lymph for adequate analysis. The sample of lymph provides an in situ molecular portrait of the lymph and the lymph-circulating tumor cells (LCTCs). We were able to dissect the critical properties of LCTCs that orchestrate their dissemination and survival in comparison to those of BCTCs from the same animal as they exit the primary tumor. We found that in contrast to BCTCs, LCTCs exist in clusters, display a hybrid epithelial/mesenchymal (E/M) phenotype and cancer stem cell-like properties and constitute extraordinarily efficient metastatic precursors. In addition, we found that EGF is the major tumor-derived factor in the lymph from metastatic tumor-bearing animals compared to nonmetastatic tumor-bearing animals and that the receptor for EGF is expressed in LCTCs but not BCTCs.

## RESULTS

### Visualization and mapping of lymphatic vessels allows the isolation of LCTCs before they reach the regional lymph nodes

We developed a novel microsurgical technique for the collection of lymph draining a primary tumor prior to its entry into the SLN. We used metastatic MTLn3 and nonmetastatic MTC cell lines that were transplanted orthotopically into the mammary fat pads of immunocompetent female Fisher 344 rats for the syngeneic model. These two cell lines were isolated from the same parent tumor, 13762NF mammary adenocarcinoma, but differed in their ability to metastasize (9). Approximately 10-14 days after injecting the cells, tumors developed in all MTLn3- and MTC-implanted rats. The PBS injection site was free of tumors (negative controls). To visualize and map the afferent lymphatic vessels prior to entering the draining SLNs to make it possible to collect the lymph, we injected approximately 10 μl of lymphazurin dye around the circumference of the primary tumor or the control injection site. After careful dissection of the skin over the tumor and lymph node area, the SLNs and afferent lymphatic vessels were identified by their green color (Fig. 1A&B) and cannulated. This procedure is similar to the procedure that is routinely performed for SLN dissection in women with breast cancer. Routinely, 20-80 μl of lymph per rat was collected before the lymph vessels collapsed. The outcome of this innovative work showed that the collection of lymph draining a primary tumor prior to its entry into the draining SLN was reliable and reproducible and yielded an adequate volume for analysis.

**Figure 1.**
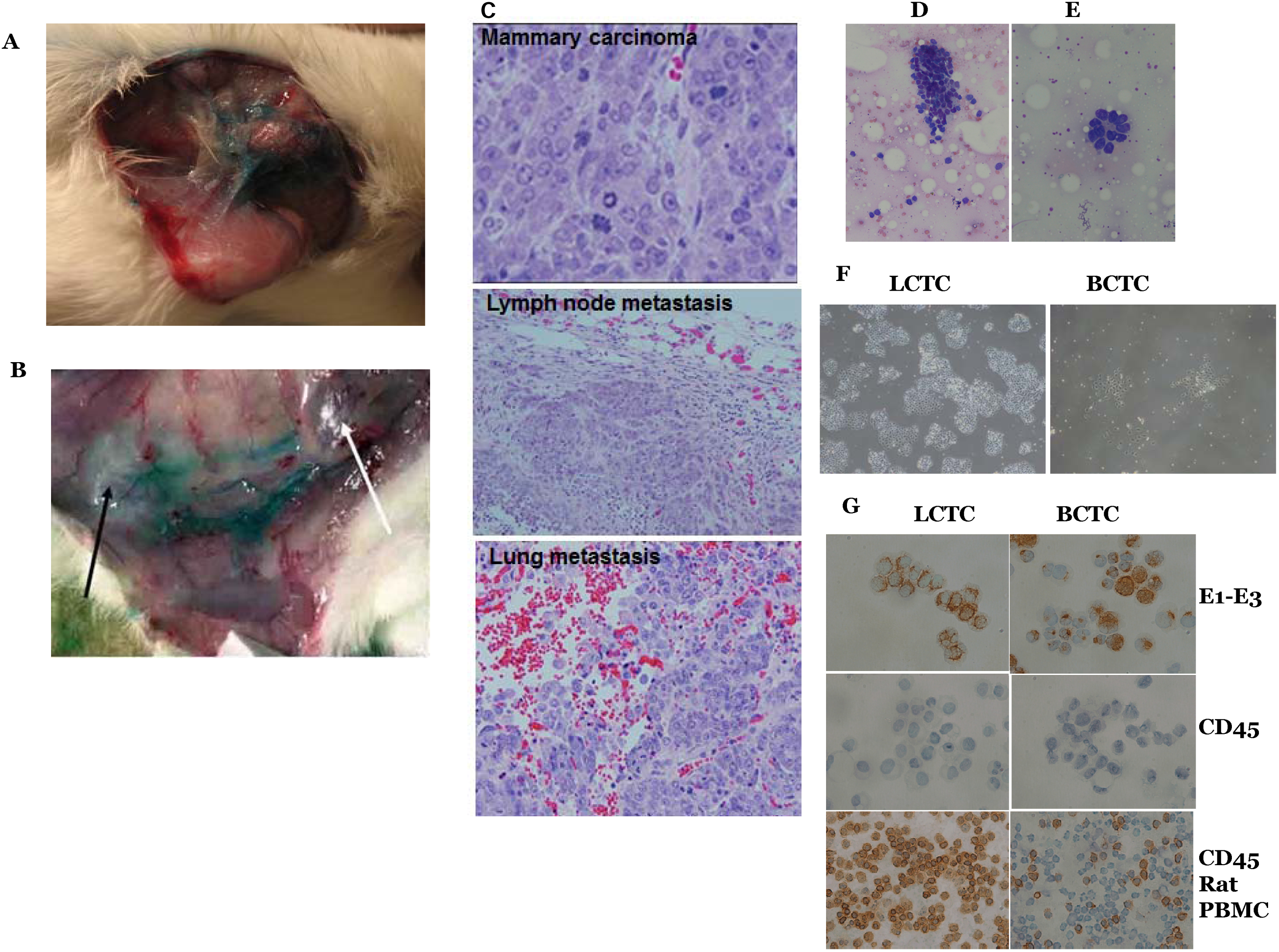
Lymph vessels visualization, lymph and LCTCs collection, LCTC identification and growth. (A) X Female rats were implanted with metastatic cell line MTLn3 and after tumor formation Lymphazurin dye was injected around circumference of the tumor and the skin over the tumor and lymph nodes area was carefully dissected away to expose the lymphatic vessels. The exposed lymphatic vessels draining from the primary tumor are shown in blue due to staining with the Lymphazurin dye. (B) The exposed primary tumor (white arrow) and the nearest lymph node (black arrow). (C) Representative images of H&E-stained histopathology showing the primary mammary tumor, and tumor cell metastases on the lymph node and lung of MTLn3 tumor-bearing rats (40X). (D) Cluster of LCTCs in directly smeared lymph on glass slides. (E) LCTCs acini in directly smeared lymph. (F) LCTCs and BCTCs growth in culture (10x). (G) LCTCs and BCTCs stained for pan-cytokeratin (AE1-AE3) and CD45. To confirm their epithelial origin. Rat white blood cells used as a control.

### Lymph node metastasis was confirmed in MTLn3 tumor-bearing animals

Histologic evaluation was used to verify the presence, number and size of metastases in the SLNs and lungs. Gross whitish colonies of tumor cells were observed in the lymph nodes and lungs of MTLn3 tumor-bearing rats but not in the lymph nodes or lungs of MTC tumor-bearing rats. The metastatic colonies were confirmed to be malignant tumor cells by histopathology (Fig. 1C).

### LCTCs existed in clusters and could be reliably harvested in the MTLn3 tumor-draining lymph prior to their entry into the SLN

In this study, we successfully identified, cannulated and collected the lymph and LCTCs on their way to the SLN. To ensure that the lymph contained tumor cells, we smeared 5 μl of the collected lymph from each animal onto a microscope glass slide for cytopathologic staining and immunohistochemistry. LCTCs were found in clumps (50- 75 cells), and a subset of LCTCs was arranged in pseudo acini (Fig. 1D & E). Blood-circulating tumor cells (BCTCs) were collected in a similar way from blood vessels (data not shown). Both cells in the lymph and the blood were cultured first in 2-dimensional (2D) monolayer cell culture plates (to separate them from white blood cells, which do not attach to the bottom of the plate or survive for a long period of time), and the attached tumor cells were washed thoroughly and then propagated in ultralow attachment cell culture plates (Fig. 1F). To confirm the epithelial origin of the cells and exclude an immune cell origin of the propagated cells, we used the accepted CTC characteristics, which include the presence of a nucleus, visible cytoplasm, and the expression of cytokeratin and the absence of CD45 expression (10), using both hematoxylin and eosin (H&E) and immunostaining. Both LCTCs and BCTCs are large in size, grow well in vitro in 3D cultures, and stain positive for cytokeratin and negative for CD45, confirming their epithelial origin (Fig. 1G). Together, these data confirmed that it is possible to collect LCTCs and BCTCs as they exit the primary tumor and that these cells can be readily propagated and identified. To avoid the effects of in vitro culture, all LCTC characterizations were performed on cells smeared directly from the lymph, and tumor cells were then collected from the slides or by using the first passage of cells before splitting.

### LCTCs and BCTCs share similar gene profiles that are distinct from those of the primary tumor and LNMs

We then determined whether LCTCs, BCTCs, primary tumors and synchronous LNMs share similar gene expression profiles indicative of the same origin. RNA was collected from cells directly from the lymph. Gene expression analysis was performed using the 770 known cancer genes from 13 canonical cancer-associated pathways that include MAPK, STS, PI3K, RAS, cell cycle, apoptosis, Hedgehog, Wnt, DNA damage control, transcriptional regulation, chromatin modification, and TGF-β in the Nanostring PanCancer Pathways Panel (Nanostring Technologies, Seattle, WA, USA). Principal component analysis was conducted to assess overall gene expression similarity across samples. LCTCs and BCTCs clustered together, while primary tumors and LNMs clustered together, suggesting that gene expression was similar between LCTCs and BCTCs and differed from that of the primary tumor and LNMs despite the same parent cell origin (MTLn3). The differentially expressed genes in LCTCs, BCTCs, and LNMs that exhibited a log2-fold change >2.0 or < −2.0 and P≤0.05 compared to primary tumors are shown in Figure 2A-C. In total, 122 and 116 genes exhibited altered expression in LCTCs vs primary tumors and BCTCs vs primary tumors, respectively. Relative to the primary tumor, LCTCs exhibited an increase in log2-fold expression for GADD55a, BAMBI, STRP4, TSPAN7, DDIT3, IL1a, and CSF3 and a decrease in log2-fold expression for COL1a2, Col5a1, COL5a2, PDGFRB, CARD11, GAQS1, IGF1, SFRP2, and COL3A1. Compared to the primary tumor, BCTCs exhibited increased expression of CSF3, WNT5B, BAMBI, MGMT, MLF1, GADD45A, HSPB1, PLAT, DUSP, and TSPAN7 and decreased expression of COL1a1, PDGFRB, COL3a1, COL5a1, IGF1, COL5a2, CARD11, GAS1, SFRP2, and COL1a2. However, only three genes were upregulated (P<0.05; >-2 log2) in BCTCs compared to LCTCs (Fig. 2D-E), given that these cells originated from the same primary tumors, with one cell type found in the blood and the other cell type found in the lymph fluid. These three genes were FBP1, plays a role in glucose metabolism, PLAT or tPA, which is a plasminogen activator that maintains blood and lymph fluidity, and FGFR2, which mediates a wide spectrum of cellular responses that are crucial for development and wound healing. Altered expression of these genes was shown to be associated with cancer progression, survival and death, and migration (11–13).

**Figure 2.**
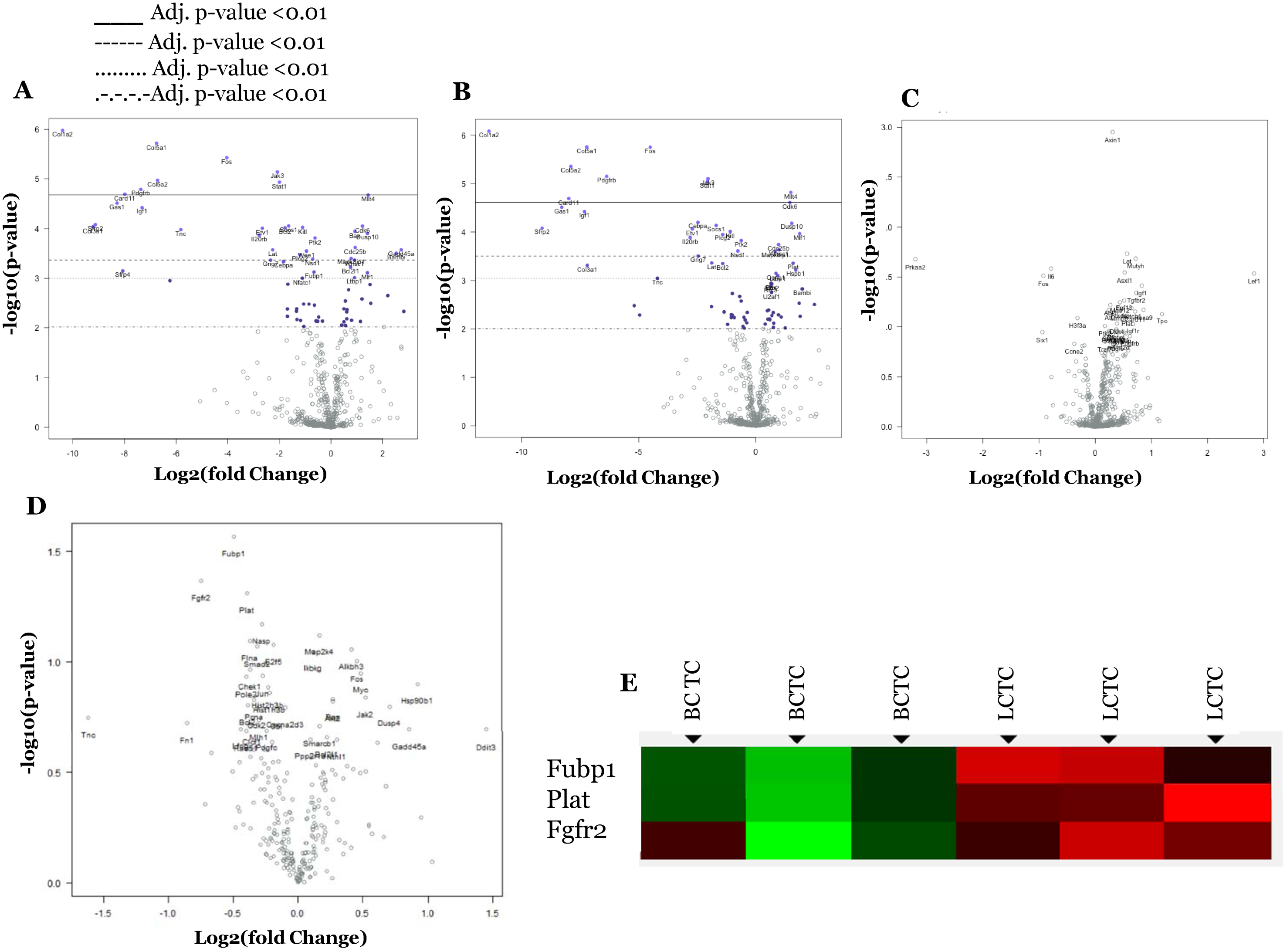
Volcano plot displaying differential expressed genes between LCTC, BCTC, LNMs (using the primary tumor as reference). The y-axis corresponds to the mean expression value of log10 (p-value), and x-axis displays the log2 fold change value. Highly statistically significant genes appear at the top of the plot above the horizontal lines (various P-value threshold indicated): p<.05, P,.01, P<.5, and highly differentially expressed genes are plotted at either side of zero. Genes were considered significant at as indicated in the figure. Only genes in significant range are colored and named. The 40 most statistically significant genes of LCTCs vs. primary tumor are shown in (A), BCTCs vs. primary tumor are shown in (B) and LMNs vs. primary tumor are shown in (C), and LCTCs vs. BCTCs are shown in (D) are labeled in the plot. (E) Differentially expressed genes between LCTC and BCTC.

To examine our differential gene expression analysis from a pathway perspective rather than the level of individual genes, we performed Pathway Score Analysis, which summarizes the data from the genes in a pathway with a single score. This approach helps in understanding which pathway scores cluster together and which samples exhibit similar pathway scores. A heatmap of pathway scores that provides a high-level overview of how the pathway scores change across samples is presented in Figure 3A. All 13 pathways examined had lower scores in BCTCs and LCTCs than in the primary tumor and LNMs (Fig. 3A). Figure 3B shows box-and-whisker plots comparing the scores of some selected pathways.

**Figure 3.**
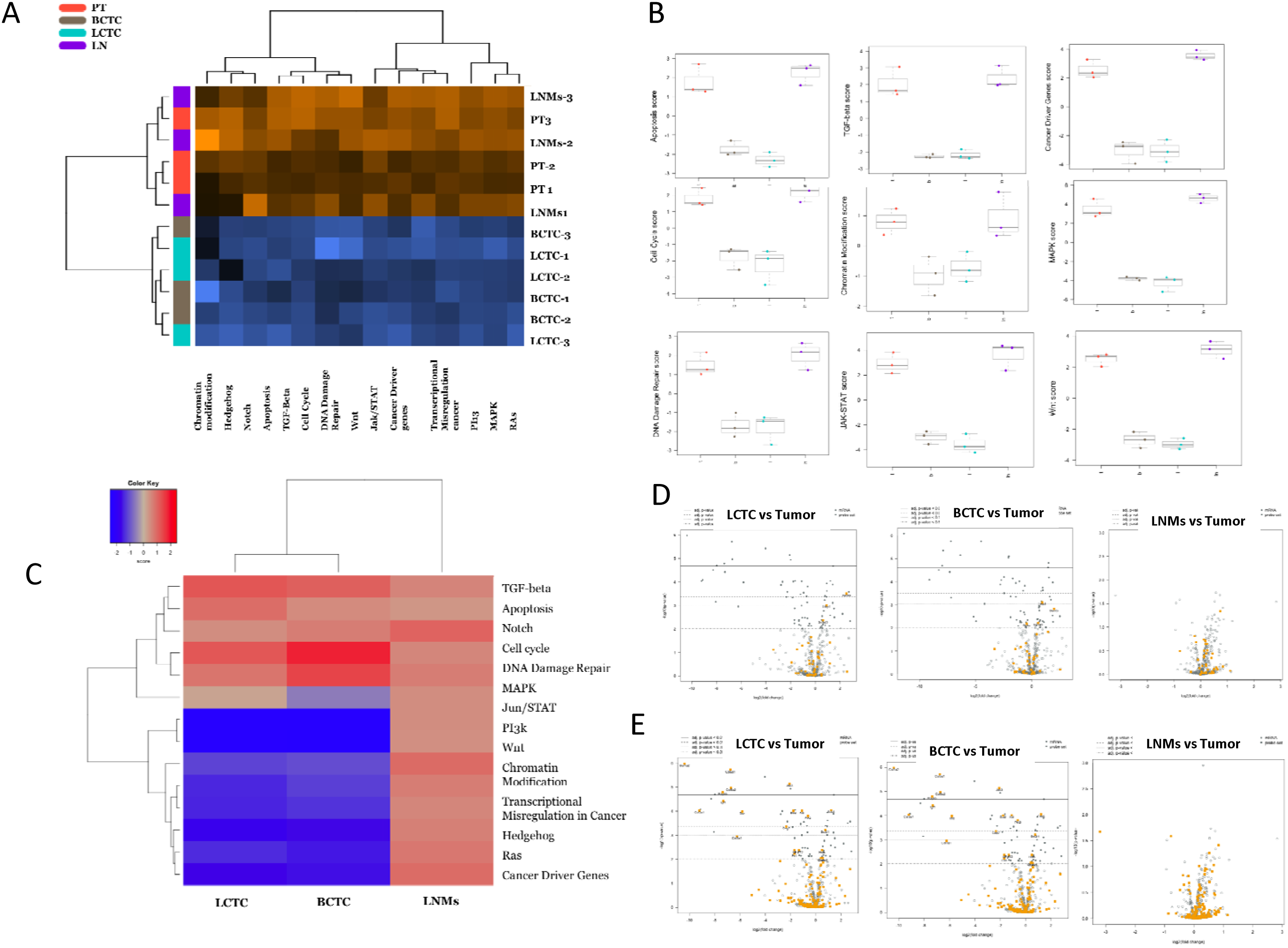
Pathway activation scores for LCTC, BCTC, LNMs, and primary tumor were compared to identify pattern of pathway activation in each. (A) LCTC and BCTC exhibit a common profile of pathway activation, while primary tumor and LNMs exhibit a common profile of pathway activation. Pathway activation score was calculated using the cumulative increase or decrease in abundance of all genes which mapped to that functional pathway. Orange indicates high scores; blue indicates low scores. Scores are displayed on the same scale via a Z-transformation (B) Box-and-whisker plots showing pathway score levels in LCTC (l), BCTC (b), primary tumor (t) and LNMs (ln). (C) Gene set analysis showing the variations in global significance scores among the gene sets in each sample. (D and E) Volcano plots displaying each gene’s -log10 (p-value) and log2 fold change for the selected covariate. (D) Showing volcano blots for TGF-beta and (E) showing volcano blots for PI3K. Highly statistically significant genes fall at the top of the plot, and highly differentially expressed genes fall to either side. Genes within the selected gene set are highlighted in orange. Horizontal lines indicate various p-value thresholds.

We then used Gene Set Analysis to assess the importance of the 13 examined canonical pathway activities in LCTCs, BCTCs, and LNMs relative to primary tumor cells. Global significance statistics were used to analyze cumulative evidence for the differential expression of genes in each pathway. Among the significantly (P<0.05) altered pathways were JAK/STAT and PI3K, which had the lowest scores in both LCTCs and BCTCs compared to primary tumors (Fig. 3C). In contrast, the TGF-β, apoptosis, cell cycle, and DNA damage repair pathways had high scores in both LCTCs and BCTCs compared with those in primary tumors (Fig. 3C). TGFβ (Fig. 3D) and PI3K (Fig. 3E) had a number of probes with significant results.

### LCTC and BCTC protein expression and phosphorylation status

To confirm the expression of the above signaling pathways at the protein level, we used the Immuno-Paired-Antibody Detection (IPAD) system provided by ActivSignal, Inc. (Natick, MA, USA). This analysis provides information on the phosphorylation states, protein levels and cleavage of more than 60 signaling factors that cover more than 20 major signaling pathways. IPAD analysis revealed the significant upregulation of proteins that affect the DNA damage response, the cell cycle, apoptosis, epithelial-mesenchymal transition (EMT), and the TGFβ and EGFR pathways in LCTCs and BCTCs (Fig. 4), confirming our gene expression data. Specifically, we show changes in cell cycle progression (the upregulation of p21 and p27 and the reduction of Rb phosphorylation) Fig. 4A), the activation of DNA damage repair (decreased phosphorylation of histone H2AX and Chk2) (fig. 4B), and the activation of apoptosis, EGFR, TGF-β (increased phosphorylation of Smad, Mek-1, and p-44) (Fig. C-E) and EMT (the upregulation of Mek-1 and p-44) (Fig. 3F). However, we observed lower expression and phosphorylation of p-27, Mek-1, and p-44 in lymph node metastases (LNMs) that in the other cell types.

**Figure 4.**
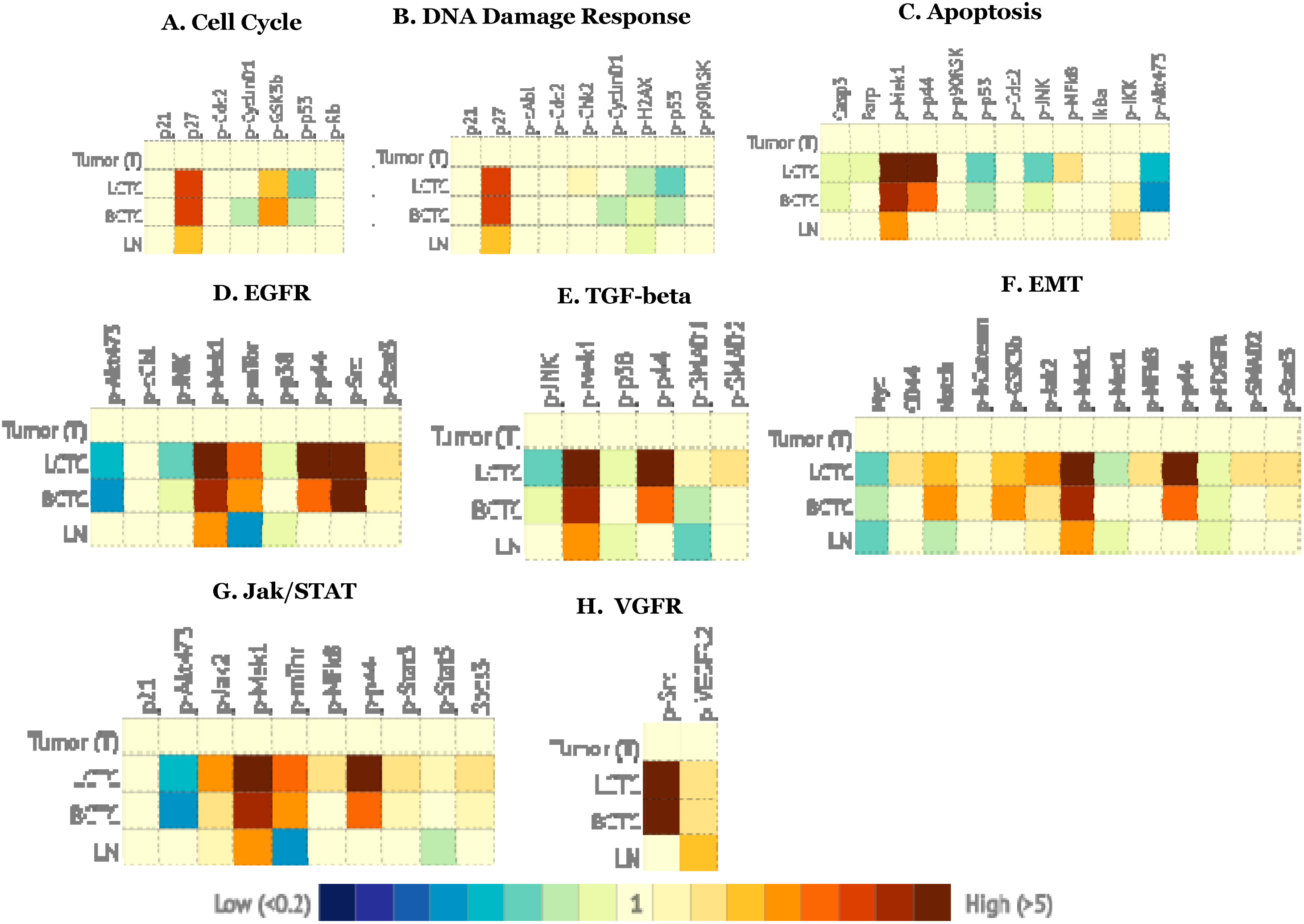
Signaling pathways alerted in LCTCs, BCTCs, and LNMs relative to primary tumor. Heat maps demonstrating changes of signaling pathways in LCTC, BCTC, and LNMs relative to primary tumor. Analysis was performed using IPAD technology by ActivSignal, Inc. The graph shows the major proteins involved in each pathway. The heat present three gradations of color intensities corresponding to 1.2, 1.8 and 2.4 and higher fold increase or decrease in IPAD values over primary tumor. Translation of the IPAD values to actual change in the activity of signaling molecules depends on the target. On average, 1.8-fold change in IPAD values corresponds to 3-fold change in the target activity. Three samples from three independent animals.

In addition, the IPAD analysis revealed other pathways that were not shown to be altered in our gene expression analysis. These pathways include NF-κB, the heat shock response and the unfolded protein response, which were similar in all samples, and signaling pathways that were upregulated only in LCTCs and BCTCs, such as mTOR, insulin receptors and IGF1R. Collectively, the gene and protein expression data indicate the upregulation of signaling pathways involved in cell motility (EGFR, EGF, ErbB2, IGF-IR, and tPA) and signaling pathways involved in angiogenesis (VEGFR and PDGFR) (Fig. 4H), cell proliferation and DNA repair.

### LCTCs display a hybrid E/M phenotype

Because our transcriptional and translational analyses showed upregulation of the TGF-β pathway in LCTCs and BCTCs, we assessed the E/M phenotypes of LCTCs and BCTCs by measuring E/M markers. During the EMT process, cells lose the epithelial markers E-cadherin and cytokeratin and gain the mesenchymal markers vimentin and N-cadherin (14). The loss of these intercellular adhesion molecules allows cells to become motile and enter the bloodstream or lymphatic system (15). BCTCs exclusively expressed the mesenchymal markers N-cadherin and vimentin, whereas LCTCs expressed vimentin and the epithelial phenotype markers E-cadherin and N-cadherin (Fig. 5A-B). LCTCs and BCTCs expressed similar levels of the EMT transcriptional factors TWIST1, ZEB1, ZEB2, SNAI1, SNAI2, and BMI1 (Fig. 5C), which organize entrance into a mesenchymal state by suppressing the expression of epithelial markers and inducing the expression of other mesenchymal markers (15). These LCTC phenotypic characteristics were consistent with the hybrid E/M state, which represents a partial or intermediate E/M phenotype (16). Cells with the hybrid E/M phenotype have both epithelial properties, such as adhesion, and mesenchymal properties, such as migration (17). These properties allow these cells to move collectively as clusters. Cells in clusters can exit the bloodstream more efficiently, are more resistant to apoptosis, and can be up to 50 times more metastatic than individually migrating cells (17). Furthermore, EMT has been associated with epithelial and carcinoma stem cell properties (18).

**Figure 5.**
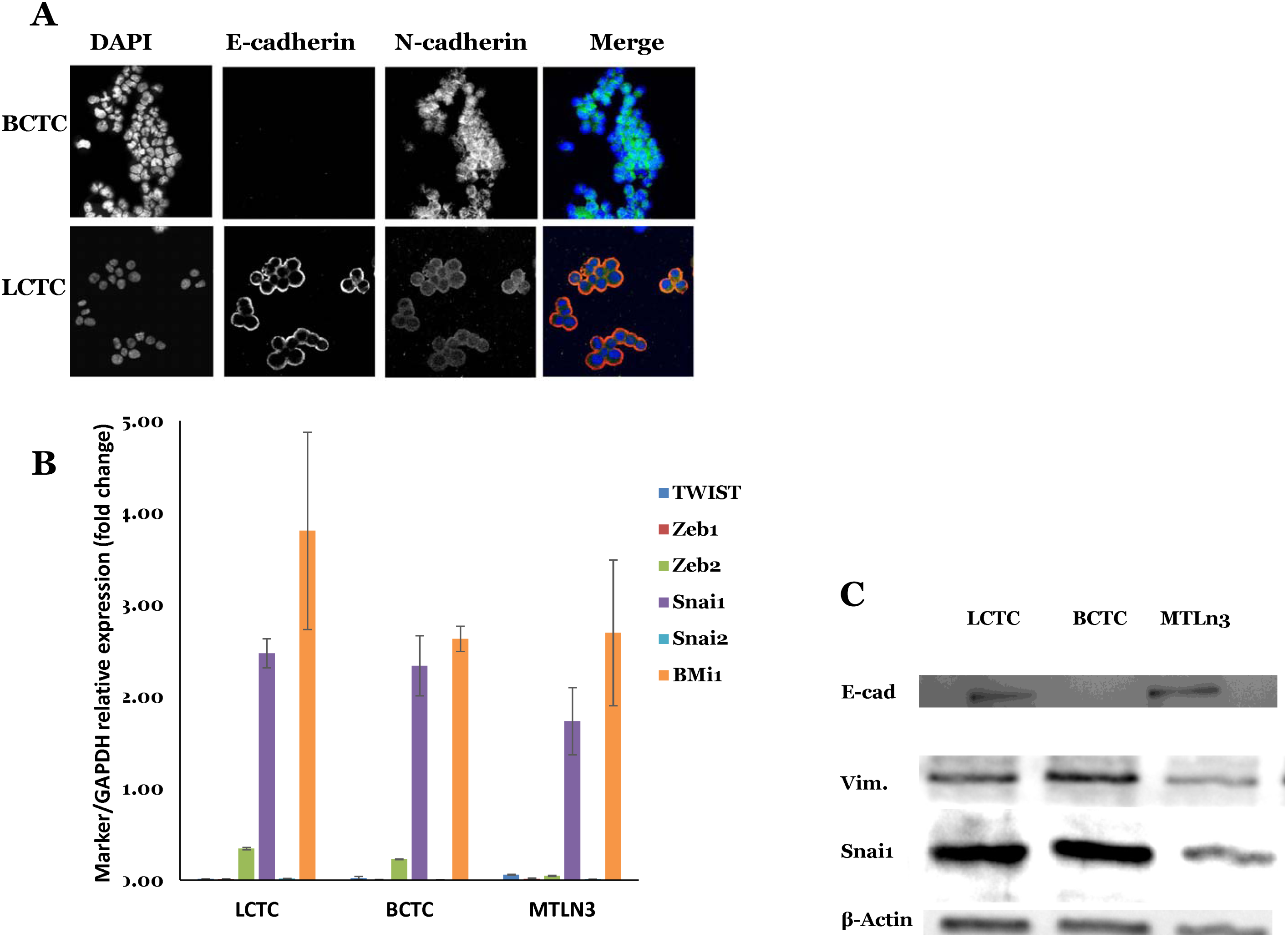
EMT phenotype and markers expressed by LCTCs, BCTCs, MTLn3. (A) EMT was recorded through immunofluorescence after cells were fixed and stained with anti-E-cadherin (red) and anti-N-cadherin (green) antibody visualization through confocal microscopy with 40X. (B) Fold change of mRNA expression relative to GAPDH of selected EMT markers (mean of three samples from different three animals, error bars denote ± SD). (C) Immunoblotting analysis of EMT markers. β-actin was used as a loading control.

### LCTCs display cancer stem cell properties and have a higher propensity than BCTCs to form mammospheres in culture and to form tumors in vivo

The induction of EMT in immortalized human mammary epithelial cells results in the acquisition of mesenchymal traits and the expression of stem cell markers, resulting in an increased ability to form mammospheres, a property associated with mammary epithelial stem cells (18). Therefore, we reasoned that cancer stem cells (CSCs) or tumor-initiating cells must be a component of LCTCs and BCTCs. Using immunohistochemistry and immunofluorescence, we examined the CSC properties of LCTCs and BCTCs using the accepted breast cancer stem markers CD29, CD44, and CD24 (19–22). LCTCs were enriched in cells that are CD29+, CD44+, and CD24+, whereas BCTCs were enriched in cells that express CD29+ and CD44+ but are CD24- or CD24low (Fig. 6A). Compared to BCTCs and MTLn3 cells, LCTCs also expressed high levels of additional CSC markers, such as NANOG, MMP9, and ALDH1 (23) (Fig. 6B, C). These data suggest that LCTCs have CSC-like properties, as demonstrated by the expression of CD29+/CD44+/CD24+ surface markers and high levels of ALDH1, which have been shown to increase in cells with stem/progenitor properties (23). By contrast, BCTCs included cells that had the phenotype CD29^+^/CD44^+^/CD24^low^. To further assess the self-renewal properties of LCTCs and BCTCs, we performed two assays that are routinely used to assess cancer cell stemness, in vitro spherical colony or mammosphere formation (24) and in vivo tumor formation in immunocompromised mice (25). Both LCTCs and BCTCs grew as nonadherent mammospheres in ultralow attachment plates (Fig. 7D); however, LCTCs formed ten times more mammospheres than BCTCs (Fig. 7D), showing the unique self-renewal ability of LCTCs (Fig. 6D). Next, we examined the tumor-initiating capacities of LCTCs, BCTCs, and the parent MTLn3 cells. To this end, we transplanted cells, after sorting them using fluorescence-activated cell sorting (FACS) in limiting dilutions, into the mammary fat pads of female immunocompetent rats. We observed that 1×106 cells was the threshold concentration for successful colonization and tumor formation for MTLn3 cells and BCTCs (Fig. 6E). BCTCs and MTLn3 cells at concentrations of 1×10^3^ or 2×10^5^ cells, respectively, failed to form tumors during the 6 weeks following implantation. In contrast, the injection of 2×10^3^, 2×10^5^, or 1×10^6^ LCTCs resulted in visible, large tumors within 2 weeks in all three rats (Fig. 6E).

**Figure 6.**
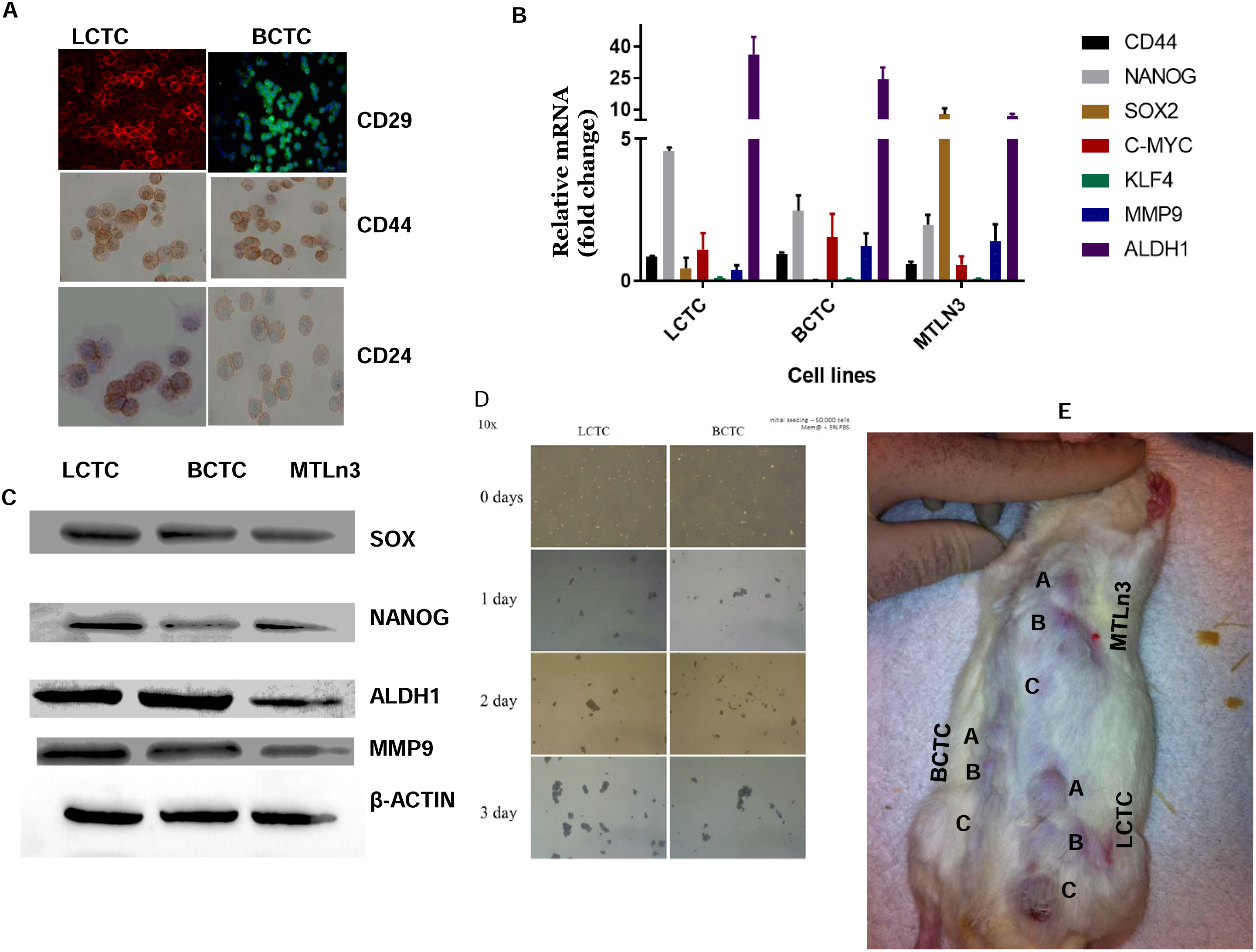
Cancer stem cell signatures of LCTCs and BCTCs. (A) Immunofluorescence analysis of CSC markers CD29 (20X), CD44 and CD24 (40X) in LCTC and BCTC. (B) RT-qPCR analysis of stem cell transcription factors mRNA levels relative to GAPDH (mean of three samples from three different animals, error bars denote ± SD). (C) Immunoblotting analysis of selected stem cell markers (D) Representative images of mammospheres at 14 days post seeding of LCTC and BCTC (magnification 20X). (E) Representative image of tumor formation in rat implanted with various numbers A=1×10^6^, B=2×10^5^, C=1×10^5^ of LCTCs, BCTCs, and MTLn3 cells. Three independent samples from three different animals. These data demonstrated that in contrast to BCTCs, LCTCs possess stem-like properties and have the ability to self-renew and efficiently form tumors.

### LCTCs and BCTCs downregulate antigen presentation pathways to escape the immune response

We next examined the immune profiles of LCTCs, BCTCs, and LNMs compared to that of the primary tumor to understand why these cells are not detected by the immune system, either in the blood or in the lymph. We performed targeted gene expression profiling using a custom 795-gene Nanostring Panel composed of immune-related genes and genes pertaining to common cancer signaling pathways (Nanostring Technologies, Seattle, WA, USA). Our undirected and directed global significance analyses (Fig. 7A) showed that the tumor necrosis factor (TNF) superfamily pathway was upregulated in LCTCs and BCTCs but not in LNMs compared with primary tumors. Although TNF is mainly produced by lymphocytes, it is also produced by tumor cells (26,27) and affects cellular processes such as apoptosis, necrosis, angiogenesis, immune cell activation, differentiation, and cell migration (28). On the other hand, the antigen presentation, pathogen response, and major histocompatibility complex (MHC) pathways were among the significantly downregulated immune pathways in LCTCs and BCTCs compared to primary tumors. These data suggest that LCTCs and BCTCs undergo immune escape and become invisible by downregulating the antigen-processing machinery. This work, for the first time, shed light on how circulating tumor cells in the lymph evade the immune system.

**Figure 7.**
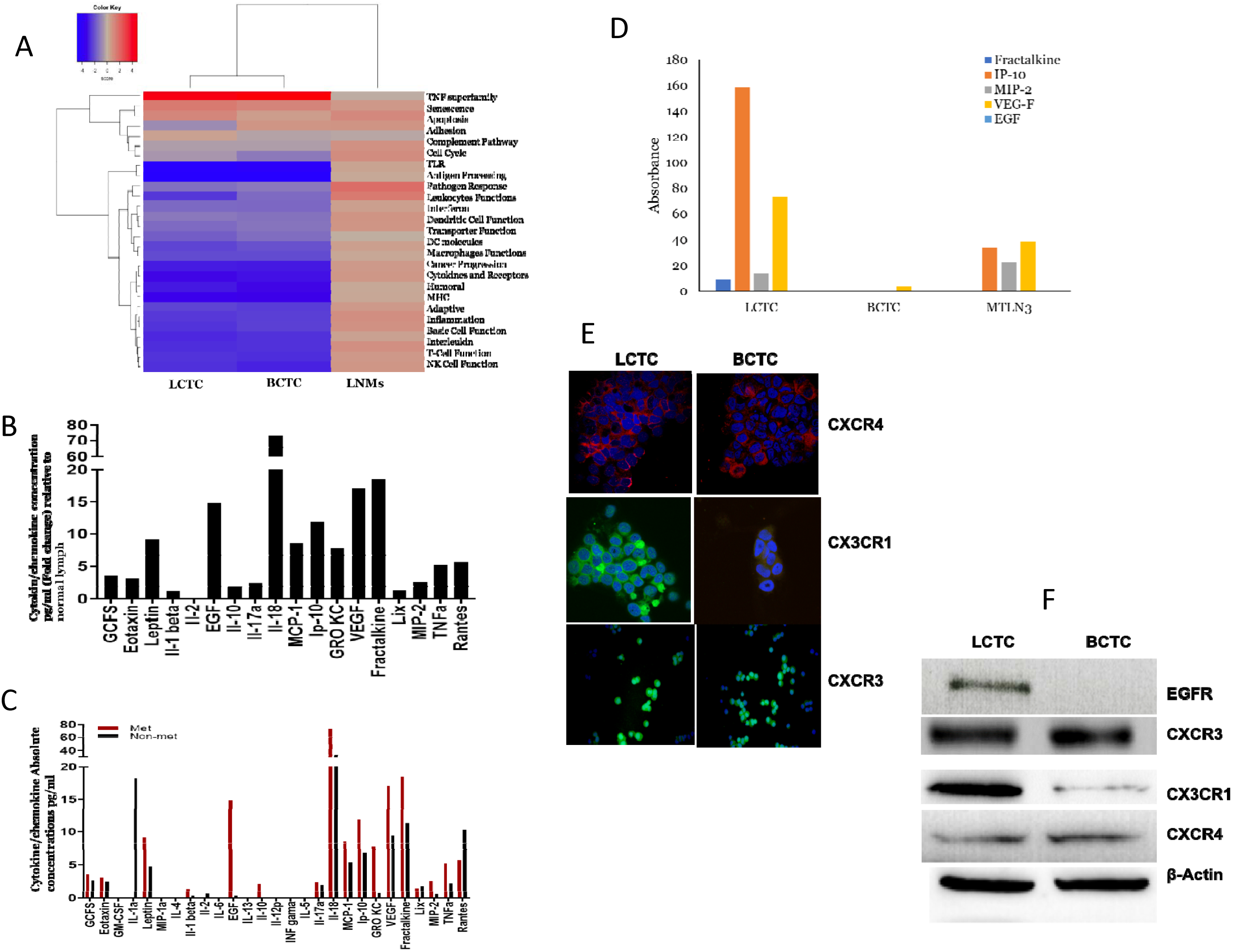
Heatmap plot of directed global significance score of immune pathway and cytokines and receptors quantification. (A) The Heatmap display the extent to which a gene set’s genes are up- or down-regulated with the variables showing the significantly expressed immune pathways in LCTCs, BCTCs, and LMNs compared to primary tumors. (B) A panel of cytokines, chemokines and growth factors measured in lymph from metastatic tumor-bearing animals relative to that of the lymph from normal animals. (C) Absolute mean concentration pg./ml of cytokines, chemokines, and growth factors in the lymph of metastatic tumor-bearing and non-metastatic tumor-bearing animals. n=three samples from three animals. (D) Absolute mean concertation pg./ml of cytokines and chemokines released by LCTCs, BCTCs, and MTLn3 cells detected in 3D culture medium supernatants. (E) CXCR4, CX3CR1, and CXCR3 expression in LCTCs and BCTCs was detected by immunofluorescent and the results were visualized by confocal microscopy. (F) Immunoblotting of EGFR, CXCR3, CX3CR1 and CXCR4. β-actin used as loading control.

### The lymph immune microenvironment

We next examined the cytokines/chemokines and growth factors in the primary tumor-draining lymph (tumor-derived factors) prior to lymph node entry to determine which factors may protect LCTCs, promote their migration to the lymph nodes and aid in premetastatic niche formation. A multiplex assay consisting of 27 cytokines/chemokines and growth factors was used to profile these factors in lymph from metastatic tumor-bearing, nonmetastatic tumor-bearing, and non-tumor-bearing control rats. Of the 27 cytokines/chemokines examined, 19 had >2-fold increases in the metastatic lymph relative to the normal lymph (Fig. 7B). EGF, TNF-α, IFN-γ-induced protein 10 (IP-10 or CXCL10), vascular endothelial growth factor (VEGF), and fractalkine (CX3CL1) showed >10-fold increases, and interleukin 18 (IL-18) showed a >70-fold increase in the metastatic lymph compared to the normal lymph (Fig. 7B). These cytokines/chemokines and growth factors play an important role in stimulating immune responses, immune cell chemotaxis, and tumor cell migration, invasion and metastasis (29–32).

Comparing the lymph from animals with tumors and metastases to that of animals with tumors but without metastases, EGF, and keratinocyte chemoattractant/human growth-regulated oncogene (GRO KC or CXCL1) showed >10- fold increases in the metastatic lymph compared to the nonmetastatic lymph (Fig. 7C). EGF is released by cells and then binds to its receptor (EGFR) on either the cell itself, stimulating its own growth, or on neighboring cells, stimulating the ability of the cells to divide (33), while CXCL1 (bind to CXCR2) plays a role in the immune response, attracts CD11b+Gr1+ myeloid cells into the tumor and enhances the survival of tumor cells facing the challenge of invading new microenvironments, tipping the balance from immune protection to tumor promotion (34,35). Moreover, IL-12p, which stimulates IFN-γ production and activates both innate (NK cells) and adaptive (cytotoxic T lymphocytes) immunity (36,37), IFN-γ, and other cytokines involved in T cell stimulation and differentiation, macrophage activation, and class II MHC expression (38,39) were not detected in the metastatic lymph in appreciable amounts relative to the those in the nonmetastatic lymph (Fig. 7C). These data suggest that the metastatic lymph microenvironment is enriched in molecules that stimulate immune responses, immune cell chemotaxis, and tumor cell migration, invasion and metastasis but is not enriched in molecules that stimulate T lymphocyte activation and gene processing and presentation, favoring the formation of an immunosuppressive microenvironment.

### Primary tumor-derived factors in the lymph are also produced by LCTCs and act in a paracrine manner

To determine if the primary tumor-derived factors produced in the lymph fluid were also produced by LCTCs and BCTCs, we performed the same multiplex assays using growth medium from LCTCs, BCTCs, and the MTLn3 cell line. The LCTCs produced >10-fold higher fractalkine (CX3CL1), IP-10 (CXCL10, which binds to CXCR3), macrophage inflammatory protein 2 (MIP-2, CXCl2/CXCl1), and VEGF levels than the BCTCs (Fig. 10A). However, none of the cells produced detectable EGF levels (Fig. 7D). Finally, we examined LCTCs and BCTCs for the receptors of these factors and found that LCTCs expressed the EGFR protein, while BCTCs did not express EGFR (Fig. 7F). Compared to BCTCs, LCTCs expressed high levels of the CX3CR1 protein (CX3CL1 receptor) but similar protein levels of CXCR3 (CXCL10 receptor) (Fig. E-F). Additionally, we compared our data for CXCR4, which is known to be expressed in peripheral blood CTCs from breast cancer patients (40). We found that both LCTCs and BCTCs expressed CXCR4 (Fig. 7E-F). These data suggest that the cytokines and chemokines found in the lymph are partly produced by LCTCs and may function in a paracrine manner. In addition, our data may provide some evidence that interactions between LCTCs and the tumor-associated lymph microenvironment could establish a potential positive-feedback loop that contributes to lymph node metastasis. This result also suggests that BCTCs differ from LCTCs and may produce different factors to survive in the blood microenvironment.

## Discussion

Analyzing the phenotypic and molecular characteristics of LCTCs and BCTCs as they exit the primary tumor and identifying the factors that orchestrate their metastatic potential is an important step for understanding the biology of these cells and the metastasis process. These characteristics may not be evident through an analysis of bulk primary or metastatic tumor populations (41) or even peripheral CTCs (P-CTCs) alone (42). P-CTCs are derived from many sources that include primary tumors, metastatic lesions in different organs, and tumor cells existing in the lymph nodes; therefore, these cells may have altered phenotypes/genotypes depending on their organ of origin (43). Here, we demonstrated that it is possible to identify the afferent lymphatic vessels and collect the lymph fluid and tumor cells therein before they reach the regional lymph nodes, as well as BCTCs as they exit the primary tumor. Thus, examining the intercellular and extracellular properties and microenvironments of tumor cells (LCTCs and BCTCs) as they exit the primary tumor in comparison to each other, the primary tumor, and LNMs may provide critical information about cancer biology and the metastatic process, which has important clinical implications.

To our knowledge, our study is the first to study tumor cells as they exit the primary tumor into the lymph en route to the lymph node. Here, we accurately identified LCTCs and BCTCs and found that LCTCs exist in clusters or clumps of 10-75 cells (Fig. 1C). Given the short distance between the tumor and the SLN and the one-way nature of cell traffic, we believe these clusters originated from the primary tumor and were not a result of multidirectional movement (44); therefore, these cells represent pure cells coming from the primary tumor and not a mixture of cells from the primary tumor and metastases, as is the case for P-CTCs. Clumps of tumor cells in the blood were initially observed by Liotta et al. (4) and were suggested to arise from oligoclonal tumor cell groupings and not from intravascular aggregation (41). We observed that LCTCs exist as clumps along the lymphatic vessels (data not shown), suggesting that they move as cohesive clusters. This observation is supported by intravital imaging studies that showed that cell clusters rather than single cells invaded through the lymphatic system instead of the blood circulation (45), suggesting that single cell motility is essential for blood-borne metastasis, while cohesive invasion is involved in lymphatic spread (45). Unlike LCTCs, large P-CTC clusters are rare in the peripheral venous circulation and constitute only approximately 2.6% of the total P-CTC population (41). P-CTC clusters have been known for many years to seed colonies with greater efficiency and were recently reported to have 50 times greater metastatic potential than individual P-CTCs (41,46). This behavior of cell clusters was reported to be due to a number of factors, including protection against anchorage-dependent apoptosis (47) and shielding from assault by immune cells (48).

We then investigated the molecular characteristics (transcriptome, proteome and immune landscapes) of these living tumor cell clusters as they exit the primary tumor en route to the lymph node and compared them to those of LNMs, the primary tumor and BCTCs. Although in our study, all tumor cells (primary tumor, LNMs, LCTCs, and BCTCs) originated from a single parent tumor cell line (MTLn3), we found striking differences in the gene expression and pathway scores of tumor cells engaged in their microenvironments (primary tumor and LNMs) and those of lymph- or blood-circulating cells (LCTCs and BCTCs). Our findings are consistent with those of studies that reported that P-CTCs are biologically different from primary tumors (49,50). The detachment of cancer cells from primary tumors and their ability to survive outside their natural extracellular matrix niches may lead these circulating cells to undergo dramatic biological changes (51,52). These findings have tremendous implications for cancer treatment because primary tumor molecular characterization currently plays an important role in the treatment strategies as well as the prognosis of breast cancer; therefore, reliance on the primary tumor characteristics can be misleading (49,50,53–55).

Our data also showed that most of the pathways examined were downregulated in LCTCs and BCTCs compared to primary tumors and LNMs, except for pathways that control DNA repair, the cell cycle, apoptosis and TGF-β. The upregulation of apoptosis, cell cycle, and DNA damage repair pathways may constitute strategies by which LCTCs and BCTCs survive stressful conditions by initiating complex signaling networks to monitor the integrity of the genome during replication and initiate cell cycle arrest, repair, or apoptotic responses if errors are detected (56). Enhanced DNA repair capabilities were reported previously in CTCs from breast cancer compared to primary tumors. This finding is important and has clinical implications, especially when treating cancer patients with DNA-damaging therapies, such as anthracyclines and platinums, which are known DNA-damaging drugs that are routinely used for breast cancer treatment (57).

Our data also showed that there were striking differences between LCTCs and BCTCs. LCTCs but not BCTCs exhibited altered TGF-β and EMT pathways and were found in clusters. One of the characteristics of these cell clusters is the coexpression of E/M markers, which is known as hybrid or partial EMT (58). In fact, LCTCs but not BCTCs exhibited a hybrid EMT phenotype, which indicates that LCTCs have mixed epithelial and mesenchymal properties, thereby allowing them to move collectively as clusters (59).

As we mentioned earlier, cells in clusters were characterized by a higher metastatic potential than cells that were not in clusters and could predict a poor prognosis in breast cancer patients (60). It was also shown that these clusters are more capable of initiating metastatic lesions than cancer cells that are moving individually with a wholly mesenchymal phenotype, having undergone complete EMT (59,60). This tumor-initiating capability is an attribute of stemness-like properties that drive metastasis and reoccurrence (25). The CSCs were shown to coexpress epithelial markers (CD24 or ALDH1) and mesenchymal markers (CD44) (22,61), as we have shown that LCTCs coexpress the CSC markers CD24, CD44, and ALDH1 (Fig. 7A), and BCTCs express only CD44 and ALDH1. This result is supported by a few recent studies that suggested that cells in a hybrid or partial EMT state are most likely than cells in a pure epithelial or pure mesenchymal state to exhibit stemness (25). Furthermore, the coexpression of both epithelial and mesenchymal genes in the same cell promotes mammosphere formation and stemness (25). Collectively, our findings showed that compared to BCTCs, LCTC clusters exhibit hybrid E/M and stemness properties and therefore constitute extraordinarily efficient metastatic precursors in breast cancer. These data comparing LCTCs to BCTCs as they exit the primary tumor allowed for the identification of a specific signature of LCTCs that provides crucial information on their stem cell properties, as well as their ability to initiate and support the formation of LNMs. More studies are needed to further elucidate the characteristics of these cells and investigate the specific molecular mechanisms involved in breast cancer progression and the development of new drugs to inhibit metastasis.

Despite the immunological power of lymph nodes, tumor cells are able to avoid immune surveillance in the lymph fluid and the lymph node, colonize the lymph node, and then migrate to distant sites. Innate and adaptive immune responses that include macrophages, natural killer cells, interferon-c (IFN-c) secretion and CD8+ cytotoxic T lymphocytes (CTLs) constitute the immunosurveillance mechanisms by which transformed cells are eliminated (62). Under this immunosurveillance mechanism, tumor cells in the lymph may develop a phenotype that helps them avoid recognition by the immune system. Consistent with this understanding, we found that LCTCs exhibit a distinct nonimmunogenic phenotype by downregulating gene processing and presentation and MHC pathways, which may significantly impair the ability of CD8+ CTLs to recognize these cells, allowing LCTCs to survive undetected despite the presence of immune cells and supporting progression and the colonization of the lymph node (63). The evasion of the immune response is a significant event in tumor development and is considered one of the hallmarks of cancer. Therefore, distinct therapeutic strategies, which depend on the biology and mechanism of immune evasion exploited by tumor cells in the lymph, may be required for restoring productive cancer immunosurveillance.

Furthermore, accumulating evidence suggests that the primary tumor releases molecules that influence the microenvironment of the SLN and make them a permissive site, known as the premetastatic niche, for receiving disseminated tumor cells and thus promoting cell proliferation and subsequent metastases (64). We reasoned that the analysis of the tumor-draining lymph may help us identify some of these factors. We showed that the metastatic lymph contains secreted factors that differ in type and expression levels from those found in the normal and nonmetastatic lymph. Specifically, the metastatic lymph had high levels of EGF, while this growth factor was not detected in the nonmetastatic lymph. Moreover, we showed that LCTCs but not BCTCs express EGFR. EGF/EGFR-induced signaling is associated with organ morphogenesis, maintenance, and repair, as well as tumor invasion and metastasis (65,66). Collectively, our data showed that the activation of EGF/EGFR signaling in the lymph and LCTCs may create a microenvironment that is conducive to metastasis, providing a rationale for efforts to inhibit EGFR signaling in lymph metastases. However, the significance of EGFR signaling in BCTCs may need to be re-evaluated.

We then assessed whether LCTCs contributed to the cytokine/chemokine pool found in lymph fluid. We showed that LCTCs released the IP-10 (CXCL10), VEGF, fractalkine (CX3CL1), and MIP-2 (CXCl2) cytokines, which were produced at high levels in the metastatic lymph (Fig. 9B). In addition, LCTCs expressed the receptors for the cytokines CX3Cl1, CXCl10, CX3CR1 and CXCR3. Our data suggest that cytokines and growth factors released by the tumor microenvironment in the lymph and LCTCs themselves may represent extracellular triggers that control the migration programs of LCTCs (67). The CX3CR1/CX3CL1 and CXCR3/CXCL10 axes have been demonstrated to be involved in the proliferation, survival and metastasis of various malignant tumor types, including breast cancer, and were suggested to predict the site of metastatic relapse (68,69). These studies support the continued examination of the CX3CR1/CX3CL1 and CXCR3/CXCL10 axes as potential therapeutic targets in patients with breast cancer.

In conclusion, we now have the capability to routinely characterize the molecular and cellular composition of tumor-derived native lymph in transit to the draining SLN. This approach will provide a new level of information that is highly relevant to our understanding of metastasis. Moreover, the contribution of LCTCs to the overall metastatic process is not fully understood, and the percentage of tumor-draining lymph cells that enter the general hematogenous circulation is unknown. The answers to these questions will provide important insights into the molecular characteristics of metastasis.

## METHODS

### Cell lines and culture condition

The cell lines used in this study were rat metastatic MTLn3 and non-metastatic MTC cells kindly provided by Dr. Segall (Albert Einstein College of Medicine, Bronx, NY). MTLn3 cell line was clonally derived from a lung metastasis of the 13762NF rat mammary adenocarcinoma (Neri et al., 1982). Both MTLn3 and MTC cell lines were cultured in Minimal Essential Medium, Alpha (MEM; Sigma, St. Louis, MO), containing nonessential amino acids (Sigma, St Louis, MO), and supplemented with 5% fetal bovine serum (FBS; Hyclone, Logan, UT). LCTCs and BCTCs were established in our laboratory from the lymph or the blood, respectively from rats with metastatic mammary tumors.

### Animal models

All experiments involving rats were conducted in accordance with National Institutes of Health regulation on the care and use of experimental animals. Purdue University Animal Use and Care Committee approved the study. Immunocompetent syngeneic female Fisher 344 rats (n=120) were purchased from Harlan (Indianapolis, IN). The rats were housed in the Purdue Animal Facility and received standard rodent chow and water ad libitum and kept at a 12-hour light-dark cycle.

### Spontaneous metastasis

To develop spontaneous metastases, rats were injected with MTLn3 or MTC cells or only PBS (vehicle control). Briefly, MTLn3 or MTC cells were grown to 70-80% confluence, trypsinized, washed with PBS and counted. 1×106 cells in 0.1 ml PBS or PBS were injected into the two left caudal- and rostral-most mammary fat pads to establish primary (MTLn3 and MTC) and metastatic tumors (MTLn3).

### Lymph fluid and blood collection

Development of the primary tumors followed by the lymph node and lung metastasis were observed after 14 days post cells implant of MTLn3 cells in rats. Tumor metastasis to the draining lymph node is grossly apparent in MTLn3-tumor bearing rats. MTLn3-tumor bearing, MTC-tumor bearing and PBS-injected animals (no tumor) were then anesthetized with Ketamine/Xylazine at 60 mg/kg of Ketamine-HCl and 5-10 mg/kg Xylazine-HCl by I.P. injections. Lymphatic vessels of tumor bearing animals and non-tumor bearing animals were visualized by injecting Lymphazurin dye (1%, isosulfan blue) (United States Surgical Corporation, Ben Venue Laboratories Inc., Ohio). Routinely, we can collect about 80-100 μl of lymph per animal. From each animal, blood was collected from blood vessels existing the primary tumor as well in addition, 3 ml of blood were collected by cardiac puncture. The primary tumor and the draining lymph node tissues were collected and processed for histopathology to confirm metastasis. Five microliters of collected lymph (80-100 μl) from each animal were immediately smeared onto a glass slide and examined under a microscope. A portion of the lymph used to grow LCTCs and another portion was used for proteomic analysis. A portion of the blood was used to grow BCTCs.

### Tumor histology and assessment of metastasis

The primary tumors, lymph nodes and lung tissues from metastatic tumor-bearing rats (implanted with MTLn3 cells), non-metastatic tumor-bearing rats (implanted with MTC cells) or the primary site of inoculation, lymph node and lung tissues from the control rats (injected only with PBS) were used for histopathological analysis. Tissues were fixed in formalin, embedded in paraffin, and 5-μm sections were stained with H&E.

### Lymph or blood circulating tumor cells isolation and propagation

To isolate and propagate the lymph or the blood circulating tumor cells, lymph (~50 μl) was mixed with Stem Cell medium EpiCult (STEMMCELL, Seattle, WA) in tissue culture dishes and incubated for 5-7 days. Plates were washed several times with PBS and a fresh stem cell medium was added. To grow cells in 3D-culture, cells were transferred to ULTRA-Low Attachment plates (Corning, Fisher Scientific, Waltham, MA USA) and were slowly adapted and cultured in Minimal Essential Medium, Alpha (MEM; Sigma, St. Louis, MO), containing nonessential amino acids (Sigma, St Louis, MO), and supplemented with 5% fetal bovine serum (FBS; Hyclone, Logan, UT). After which their epithelial nature were determined by staining with cytokeratin (AE1/AE3+8/18), CD45 (BD pharmaingen (554875) from Biocare (Pacheco, CA) to exclude the white blood cells using the rat while blood cells as a positive control (purified from the same rat blood cells using Ficoll gradient) and a negative control without primary antibody.

### ActivSignal IPAD assay

The cells from lymph or allowed to grow to have enough cell number (first oassage) were collected and lysed in PBS + 1%NP40 lysis buffer. The lysates were sent to ActivSignal for further processing (http://www.activsignal.com). ActivSignal IPAD platform is a proprietary technology for analyzing the activity of multiple signaling pathways in one reaction. Activities of more than 20 signaling pathways are monitored simultaneously in a single well through assessing expression or protein phosphorylation of 70 target human proteins. The technology allows detection of targets with high specificity and sensitivity due to combination of two distinct antibodies per each target. Each pathway is covered by multiple targets.

### NanoString nCounter analysis

RNA from cells and tissues was harvested using a RNA Isolation Kit (Roche) as per the manufacturer’s instructions. All RNA was quantified used the DeNovix DS-11 Spectrophotometer. Samples were processed for analysis on the NanoString nCounter Flex system using the 770 gene PanCancer Pathways Plus panel (606 critical genes from 13 canonical cancer pathways, 124 cancer driver genes, and 40 reference genes) and nCounter PanCancer Immune Profiling Panel from NanoString Technologies (Seattle, WA, USA), as per manufacturer’s instructions.

### RNA expression analysis

Resource complier (RCC) data files were imported into NanoString nSolver 3.0 and further analyzed using the PanCancer Pathways Advanced Analysis Module, which normalizes gene expression to a set of positive and negative controls genes built into the platform. Using the nCounter Analysis software, we identified a list of genes with significantly altered expression between LCTC, BCTC, LMNs and primary tumors. The fold change and P values were calculated using nCounter default settings. As recommended, genes whose expression levels were at or below the level of the negative controls were removed from analysis. With the remaining list of genes on the PanCancer panel, a filter cutoff of fold change ≥ ±1.5 or ≥ ±2 and P value < 0.05 were used to identify the significant gene expression changes based on the nCounter analysis. A pathway score was calculated using nSolver Advanced Analysis from the expression levels of the relevant genes in 13 canonical pathways using measurements of pathway activity values derived from singular value decompositions. This method uses metagenes to represent pathway activity and aims to capture not only over-represented significantly altered genes but also smaller but cumulatively impactful changes within a pathway.

### Lymph chemokines/cytokines determination

Chemokines/cytokines levels in the lymph collected from metastatic tumor-bearing rats, non-metastatic tumor-bearing rats and normal control rats (total n=30) and supernatant from cells cultures were measured using MILLIPLEX Rat Expanded Cytokine/Chemokine MAGNETIC Bead Premixed 27 Immunology Multiplex Assay (EMD Millip0re, Billerica, Massachusetts, USA) as per the manufacturer’s instructions. Standard curves were generated from known concentrations of each cytokine then used to determine the quantity of cytokine in each sample based on the level of spectrophotometric absorbance of the sample using regression analysis. Each assay was performed in duplicate, and each value shown in the figures is the mean of the duplicates.

### Quantification of Real-time PCR

Total RNAs were extracted from LCTCs, BCTCs, using TRIzol RNA Isolation Reagents as per the manufacturer’s instructions (Invitrogen, Carlsbad, CA). RNAs were reverse-transcribed by oligo(dT) primer using Superscript RT-PCR kit from Roche (Basel, Switzerland), according to the manufacturer’s instructions. PCR reaction was performed under the following conditions: 940C for 3 min; 94°C 30 Sec; 58°C for 30Sec; 72°C for 30 Sec for 40 cycles; and 72°C for 10min, using IQ SYBR Green Supermix Kit from Roche. Results were analyzed by the relative quantification method and expressed as relative RNA levels (∆CT, difference of cycling threshold). ∆CT values represent CT [gene]-CT [GAPDH], thus higher values indicate relatively lower expression levels.

### Western Blot analysis

Cells were grown in 3D-cultures and proteins were isolated using RIPA buffer (0.5M Tris-HCl, pH 7.4, 1.5M NaCl, 2.5% deoxycholic acid, 10% NP-40, 10mM EDTA). Protein concentrations were determined using Pierce BCA Protein Assay Kit according to the manufacturer’s instructions (Thermo Fisher Scientific). Ten micrograms of each sample protein was subjected to SDS-PAGE, and transferred to nitrocellulose paper. The blots were reacted sequentially with primary antibodies; HRP-conjugated goat anti-rabbit IgG or goat anti-mouse IgG, and visualized with diaminobenzidine.

### Immunocytochemistry

Cells were fixed with 1:1 methanol: Acetone, and preblocked with 1% bovine serum albumin in PBS. Cells were then incubated with the following anti-primary antibodies E-cadherin, N-cadherin, CX3R3, CXCR4, CXCR4 (Novus, Littleton, CO) at 4°C overnight, followed by the secondary antibody conjugated with Alexa Fluor 594 or FITC (Jackson ImmunoResearch, West Grove, PA). The cells were mounted with mounting medium containing 1 μg/ml DAPI (4′,6′-diamidino-2-phenylindole; Sigma, St Louis, MO).

### Statistical analysis

All data are presented as means ± standard deviation (SD). Statistical calculations were performed with Microsoft Excel analysis tools. Differences between individual groups were analyzed by paired t test. P values of <0.05 were considered statistically significant.

## ACKNOWLEDGMENTS

We thank Dr. Chun-Ju Chang and Mi Ran Kim for their help with EMT confocal image and Dr. Segall from Albert Einstein College of Medicine, Bronx, NY for providing the MTLn3 and MTC cell lines.

